# On the Apportionment of Population Structure

**DOI:** 10.1101/033852

**Authors:** Yaron Granot, Omri Tal, Saharon Rosset, Karl Skorecki

**Affiliations:** Rappaport Faculty of Medicine and Research Institute, Technion-Israel Institute of Technology, and Rambam Medical Center, Haifa, Israel; Max Planck Institute for Mathematics in the Sciences, Inselstr. 22-26, 04103 Leipzig, Germany; School of Mathematical Sciences Tel Aviv University, Tel Aviv, Israel

## Abstract

Measures of population differentiation, such as F_ST_, are traditionally derived from the partition of diversity within and between populations. However, the emergence of population clusters from multilocus analysis is a function of genetic *structure* (departures from panmixia) rather than of diversity. If the populations are close to panmixia, slight differences between the mean pairwise distance within and between populations (low F_ST_) can manifest as strong separation between the populations, thus population clusters are often evident even when the vast majority of diversity is partitioned within populations rather than between them. For any given F_ST_ value, clusters can be tighter (more panmictic) or looser (more stratified), and in this respect higher F_ST_ does not always imply stronger differentiation. In this study we propose a measure for the partition of structure, denoted E_ST_, which is more consistent with results from clustering schemes. Crucially, our measure is based on a statistic of the data that is a good measure of internal structure, mimicking the information extracted by unsupervised clustering or dimensionality reduction schemes. To assess the utility of our metric, we ranked various human (HGDP) population pairs based on F_ST_ and E_ST_ and found substantial differences in ranking order. In some cases examined, most notably among isolated Amazonian tribes, E_ST_ ranking seems more consistent with demographic, phylogeographic and linguistic measures of classification compared to F_ST_. Thus, E_ST_ may at times outperform F_ST_ in identifying evolutionary significant differentiation.

## Introduction

Genetic differentiation among populations is typically derived from the ratio of within-to between-population diversity. The most commonly used metric, F_ST_, was originally introduced as a fixation index at a single biallelic locus [1], and subsequently adapted as a measure of population subdivision by averaging over multiple loci [2–3]. F_ST_ can be expressed mathematically in terms of population diversities as *F_ST_*=*1−S/T*, where S and T represent the heterozygosity in subpopulations and in the total population, respectively [4–5]. The validity of F_ST_ as a measure of differentiation has been brought into question, especially when gene diversity is high (e.g., in microsatellites), and various metrics, including G’_ST_[6] and Jost’s*D* [7], have been proposed to address this inadequacy (though see [8] for a counter-perspective).

Although these metrics vary considerably in their formulation, they all follow the same basic framework of partitioning genetic diversity into within-vs. between-group components. It has long been noted, however, that the apportionment of diversity [9] does not directly reflect the strength of separation between populations, and the emergence of population clusters has been demonstrated both empirically [10] and mathematically [5, 11–12] even when the vast majority of diversity is within rather than between populations. For example, humans sampled from across Europe [13] form identifiable clusters with pairwise F_ST_ as low as 0.001, even though 99.9% of the variation is contained within populations and only 0.1% is between them. Clearly, these clusters reflect an aspect of population differentiation that is not directly captured by F_ST_, yet there is currently no commonly used metric for partitioning structure into within-and between-population components in the same way that F_ST_ partitions diversity. Dimensionality reduction schemes such as principal component analysis (PCA) [14] and clustering algorithms such as the widely used STRUCTURE [15] are highly popular, however such programs are primarily used for visualization, and there is still value in summary statistics for quantifying complex datasets on a simple 0-1 scale.

Here we propose a novel measure, denoted E_ST_, which is based on the variation in pairwise genetic distance (which we show to be a measure of internal structure), thus exposing the excess structure inherent in the total population compared to subpopulations. Conceptually, E_ST_ is formulated in three steps: 1. Population structure is defined in terms of departures from *panmixia*. 2 Panmixia is defined in terms of pairwise genetic *equidistance* between individuals (we show that a population is panmictic if all individuals are equally distant from each other). 3. Departures from equidistance are defined in terms of the *standard deviation* of pairwise distances. E_ST_ reflects the decrease in panmixia when subpopulations are pooled. The basic formula is E_ST_=1-SD_S_/SD_T_, where SD_S_ and SD_T_ represent the standard deviations of pairwise distances in subpopulations and in the total population respectively. While F_ST_ is weighed down (diminished) by high *diversity* within populations, E_ST_ is weighed down(diminished) by high *diversity* within populations, E_ST_ is weighed down by high *structure* within populations.

A core insight of this proposal is that the asymptotic (in terms of number of SNP loci considered) standard deviation of pairwise genetic distances is a good unsupervised measure of internal structure, a statistic that mimics the information extracted by dimensionality reduction and clustering schemes, thus justifiable as a basis for the definition of E_ST_. In particular, we show (in Appendix A) that a population is panmictic if and only if all individuals are *asymptotically* equally distant from each other, and that the standard deviation of pairwise genetic distances is highly associated with the deviation from panmixia and faithfully reflects the structure extracted by PCA.

## Results and Discussion

### Partitioning Diversity vs. Partitioning Structure

The difference between partitioning diversity and structure within and between two populations from the human genome diversity project (HGDP) [16] is illustrated in Figure 1. By zooming into a Russian and Chinese neighbor-joining tree of individual similarities we observe three layers of diversity and structure. Distances between individuals (black) account for most of the diversity, followed by the between-population component (red) and lastly, structure within populations (blue). The striking symmetry in the full-sized tree (1A) suggests high levels of panmixia in these two populations. Even at 100x magnification, most intra population branches (blue) are shorter than the 1x individual branches (black), indicating that these two sample sets are >99% panmictic. The red/black branch length ratio can be perceived as a rough proxy to the fixation index F_ST_ and (one minus) the blue/red branch length ratio can be perceived as a rough proxy to the equidistance index E_ST_.

**Figure 1.**
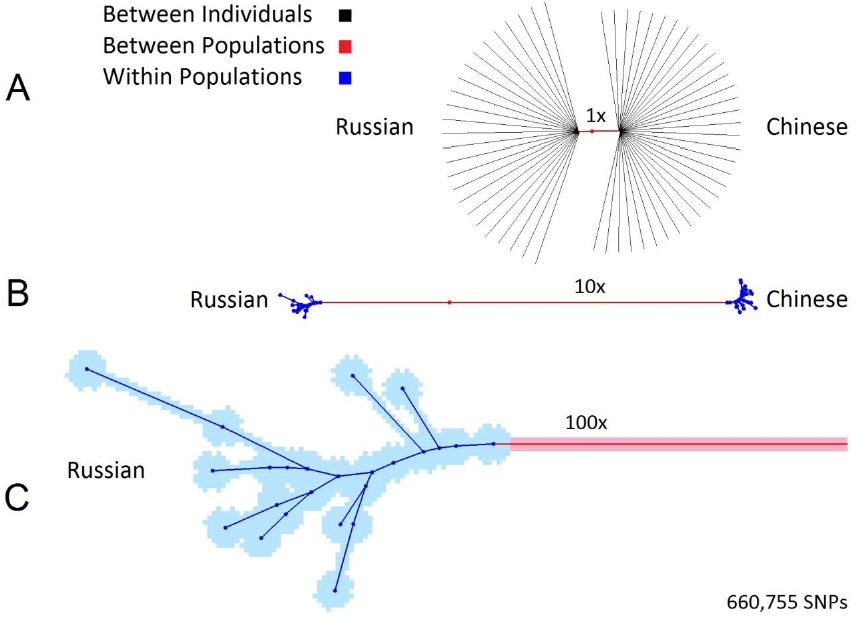
Zooming into a Russian (n=25) and Chinese (n=34) Neighbor Joining tree of individual similarities. (A) The length of the red branch compared to the overall tree length is a rough proxy to F_ST_. (B) 10x magnification highlights the structure within and between populations. The 1-blue/red mean line length is a rough proxy for E_ST_. (C) 100x magnification reveals fine substructure within Russian samples. Individual branches (black) were removed in B and C to reduce clutter.

We compared F_ST_, E_ST_, and clustering among Russian and Chinese samples, with an increasing amount of single nucleotide polymorphisms (SNPs) ranging from 10 to 660,755 (Figure 2). Using multidimensional scaling (MDS), the two population clusters gradually diverge as SNP count increases, with no corresponding increase in F_ST_. At the same time we observe a steady increase in E_ST_ directly corresponding to the emerging clusters, indicating that the Russian and Chinese HGDP samples are close to panmixia. With few SNPs this is obfuscated by the variance of the genetic distance measure, hence E_ST_ is relatively small. The actual levels of panmixia become increasingly evident as more SNPs are added, thus revealing the population clusters [11]. However this process does not proceed indefinitely; the finite number of pairwise differences among humans (~3 million SNPs) sets an upper limit to the number of available markers, and the amount of extractable information is further reduced by physical linkage. In our HGDP data the increase in E_ST_ as a function of marker count reaches a plateau approximately above 100,000 SNPs (Figure 3). Although this upper bound can vary across different datasets and types of markers, it suggests that resolution may not improve substantially with further increases in marker count. Thus, these clusters can be considered close approximations of the “true” strength of separation among these populations. For this reason, when estimating E_ST_ one should include as many markers as possible, although at a certain point additional markers provide greatly diminishing returns.

**Figure 2.**
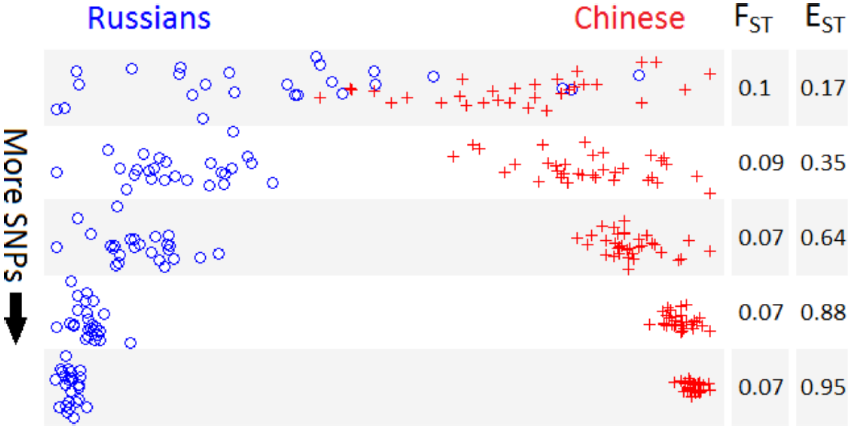
F_ST_ and E_ST_ vs. Clustering with increasing SNP count. Multidimensional scaling (MDS) plots with Russian (n=25) and Chinese (n=34) samples with increasing SNP count from top to bottom (10, 100, 1000, 10,000, and 660,755 SNPs). Two clusters gradually emerge as SNP count increases, along with an increase in E_ST_, while F_ST_ remains relatively constant.

**Figure 3.**
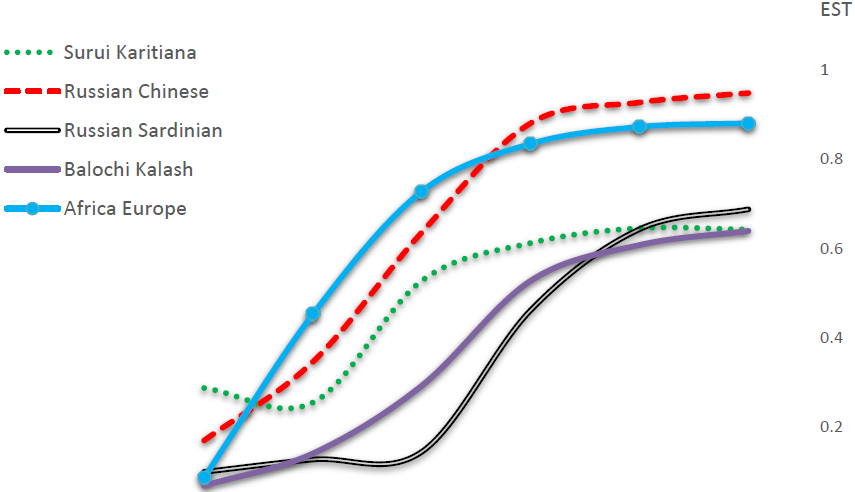
E_ST_ as a function of SNP sample size. E_ST_ was estimated in various population pairs with gradually increasing SNP sample size from 10 to 660,755. As expected, E_ST_ initially rises rather steeply, but tends to plateau before reaching the 660,755 SNP point. This suggests that we are approaching the maximal resolving power of genetic markers in this dataset, and adding markers beyond this point should not have a significant effect on -cluster separation and E_ST_.

In order to determine whether or not E_ST_ adds insight to the analysis of population structure, we sought to compare the rank order of population differentiation using F_ST_ and E_ST_. Pairwise F_ST_ and E_ST_ values from various HGDP populations are given in Table 1 (see Table S1 and Figure S1 for additional comparisons). It is noted that for almost all population pairs E_ST_ is larger than F_ST_, and only the Colombian-Maya pair entails a slightly lower E_ST_ than F_ST_, presumably due to a combination of relatively low differentiation and high levels of intra-population structure. According to the HGDP browser (http://spsmart.cesga.es/search.php?dataSet=cephstanford) the Colombians (n=7) are the only HGDP population sample where two different tribes (Piapoco and Curripaco) were combined, which can help explain the high level of structure observed in this particular population (see Table S1, Figure S9, and Materials and Methods for further analysis of E_ST_ range). There is a moderate positive correlation (r=0.61) between F_ST_ and E_ST_ among all 60 population pairs included in our analysis (Figure 4), which is consistent with the two measures capturing somewhat different aspects of population structure.

**Table 1.**
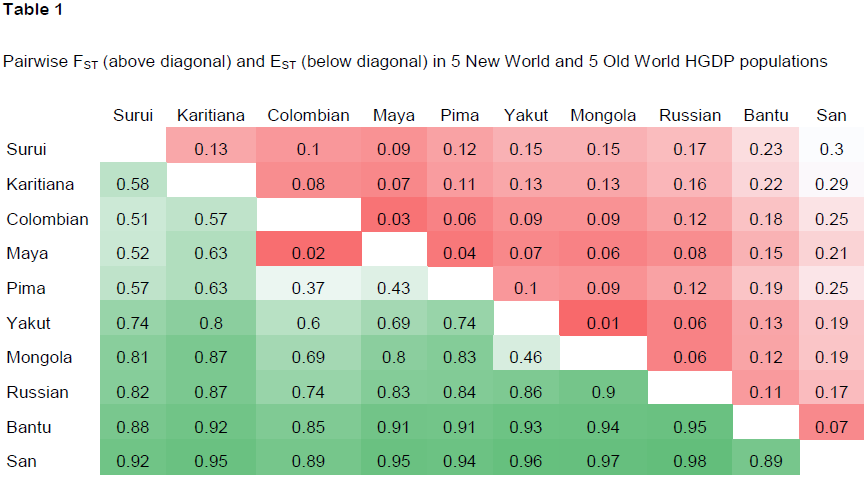
Pairwise F_ST_ (above diagonal) and E_ST_ (below diagonal) in 5 New World and 5 Old World HGDP populations

|  | Surui | Karitiana | Colombian | Maya | Pima | Yakut | Mongola | Russian | Bantu | San |
| --- | --- | --- | --- | --- | --- | --- | --- | --- | --- | --- |
| Surui |  | 0.13 | 0.1 | 0.09 | 0.12 | 0.15 | 0.15 | 0.17 | 0.23 | 0.3 |
| Karitiana | 0.58 |  | 0.08 | 0.07 | 0.11 | 0.13 | 0.13 | 0.16 | 0.22 | 0.29 |
| Colombian | 0.51 | 0.57 |  | 0.03 | 0.06 | 0.09 | 0.09 | 0.12 | 0.18 | 0.25 |
| Maya | 0.52 | 0.63 | 0.02 |  | 0.04 | 0.07 | 0.06 | 0.08 | 0.15 | 0.21 |
| Pima | 0.57 | 0.63 | 0.37 | 0.43 |  | 0.1 | 0.09 | 0.12 | 0.19 | 0.25 |
| Yakut | 0.74 | 0.8 | 0.6 | 0.69 | 0.74 |  | 0.01 | 0.06 | 0.13 | 0.19 |
| Mongola | 0.81 | 0.87 | 0.69 | 0.8 | 0.83 | 0.46 |  | 0.06 | 0.12 | 0.19 |
| Russian | 0.82 | 0.87 | 0.74 | 0.83 | 0.84 | 0.86 | 0.9 |  | 0.11 | 0.17 |
| Bantu | 0.88 | 0.92 | 0.85 | 0.91 | 0.91 | 0.93 | 0.94 | 0.95 |  | 0.07 |
| San | 0.92 | 0.95 | 0.89 | 0.95 | 0.94 | 0.96 | 0.97 | 0.98 | 0.89 |  |

**Figure 4.**
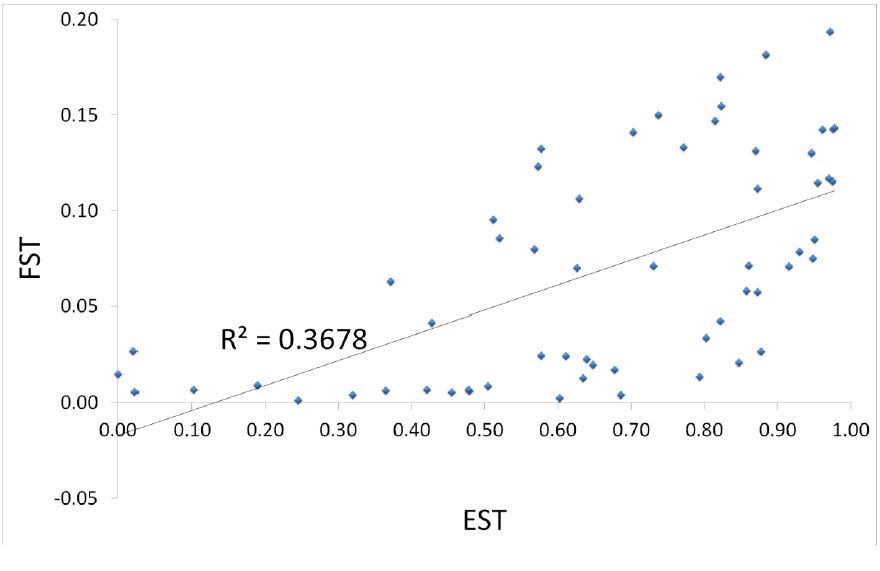
Positive correlation between F_ST_ and E_ST_. R2 = 0.3678 and r = 0.61. Note that E_ST_ has a much broader range, spanning nearly the entire 0-1 interval while F_ST_ only goes as high as 0.2 in these HGDP populations.

### Amazonians vs. Global Populations

A prominent example of the divergent behavior of E_ST_ and F_ST_ is the Surui and Karitiana, which have an unusually high pairwise F_ST_. In fact, the Karitiana are as diverged from the neighboring Surui in terms of F_ST_ as they are from the Mongola on the other side of the world (Table 1, Figure 5, and Figure S5). F_ST_ decreases initially with distance from the Amazon, from 0.13 between the two Amazonian tribes, to 0.08-0.1 between Amazonians and Colombians, and further decreases to 0.07-0.09 between Amazonians and the more distant Maya. Remarkably, the highest F_ST_ among all HGDP Native American populations is between the two geographically closest populations, the Surui and Karitiana. These apparent anomalies can be explained by the inflation of F_ST_ among genetic isolates. F_ST_ between pairs of isolates can be nearly twice as high as between either one of the isolates and a more cosmopolitan population, as pairwise F_ST_ reflects the *combined* isolation of both populations. Since the Surui and Karitiana are both isolated, their pairwise F_ST_ is nearly double that between any one of them and a larger, less isolated population such as the Maya.

**Figure 5.**
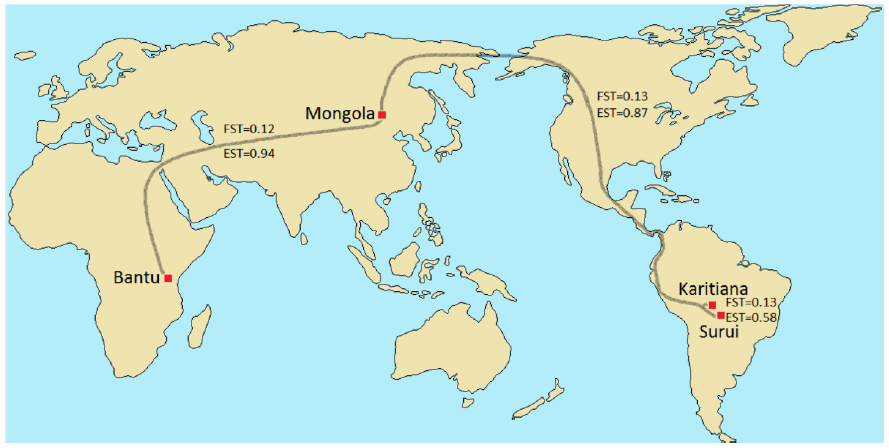
Geographic distance vs. F_ST_ and E_ST_ in various populations. In terms of F_ST_, the Karitiana are roughly as diverged from the nearby Surui (F_ST_=0.13) as they are from the Mongola on the other side of the world (F_ST_=0.13) or as the Bantu are from the Mongola (F_ST_=0.12). In terms of E_ST_, differentiation is far greater among these global populations (E_ST_≈0.9) than between the neighboring Amazonian tribes (E_ST_≈0.6).

Differentiation based on E_ST_ (Surui-Karitiana=0.58, Karitiana-Mongola=0.87, and Mongola-Bantu=0.94) seems more consistent with the geographic distances among these populations (Figure 5). It should be noted that the Surui-Karitiana E_ST_ might be somewhat underestimated due to cryptic sampling of close relatives [17], however the wide range of heterozygosity values (which are less sensitive to the sampling of close relatives) and the elevated structure across all Native American HGDP populations (Figures S3-S5) suggest that this is not merely a sampling artifact. In some cases E_ST_ also decreases with distance from the Amazon (Table 1), however this decrease is more moderate than the decrease in F_ST_ (Figure S5).

Neighbor-joining trees of individual similarities [18] are a convenient tool for representing multidimensional genetic data on a two-dimensional plane, while simultaneously displaying distances within and between populations. Two pairs of such trees, for Surui-Karitiana and Yoruba-Russians, are given in Figure 6, and we can see that in both cases distances are greater between individuals (black branches) than between populations (red branches) (Figure 6A).

**Figure 6.**
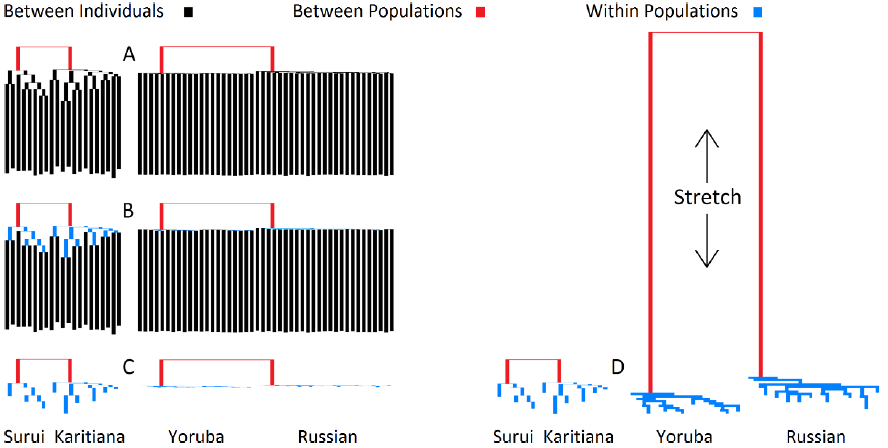
Surui-Karitiana vs. Yoruba-Russian NJ trees of individual similarities. (A) Diversity is apportioned into individual (black) and population (red) components. (B) A third component, structure within populations (blue), is added. (C) The individual component is removed. (D) The Yoruba-Russian tree is stretched to roughly match the level of structure within the Surui-Karitiana tree.

The ratio of within- to between-population distance is roughly equivalent in the two population pairs, however the Yoruba-Russian tree is significantly *flatter*, indicating greater panmixia within these two populations (Figures S6-S7). Adding a third dimension of intra-population structure (blue branches) highlights this discrepancy (Figure 6*B*), which is further accentuated by removing the inter-individual component (Figure 6*C*) and stretching the Yoruba-Russian tree to match the level of structure observed in the Surui-Karitiana tree (Figure 6*D*). At first glance the Amazonian tribes, with their long population branches, appear to be as differentiated as the Yoruba are from the Russians. Upon closer inspection, however, the Yoruba and Russians appear more strongly diverged. The Amazonian tribes are highly structured not only between them, but also within them, resulting in distant, but loosely separated clusters. This aspect of population structure is not captured by F_ST_, which is actually slightly higher between the Surui and Karitiana (0.13) than between Yoruba and Russians (0.12), but is revealed by the higher E_ST_ between Yoruba and Russians (0.97) compared to the Surui and Karitiana (0.58).

### E_ST_ and the Dissimilarity Fraction

Witherspoon et al. [10] have also examined population structure through the lens of pairwise genetic similarities and dissimilarities. They have defined the dissimilarity fraction, ω, as the probability that individuals are genetically more similar to members of a different population than to members of their own population. An intuitive proxy for ω is (half) the overlap of the within and between pairwise distance distributions. For population pairs, this probability has a 0-0.5 range, with the extremities ω=0 indicating that individuals are always more similar to members of their own population and ω=0.5 indicating that individuals are just as likely to be more similar to members of the other population as to members of their own population (see [5] for a formal analysis of such a metric and its relation to classification accuracy). Witherspoon et al. reported that that when many thousands of loci are analyzed, individuals from “geographically separated populations” are never more similar to each other than they are to members of their own populations. The definition of “geographically separated” is, of course, open to interpretation. We found no overlap (ω=0) between the Adygei and Uygur HGDP samples, but some overlap (ω>0) between Mayans and Surui, despite a 4x higher F_ST_ (Figure 7). Thus, F_ST_ and the dissimilarity fraction (ω) are not necessarily congruent. The E_ST_ values for these two population pairs are more consistent with ω, showing strong separation between the Adygei and Uygur (0.79) and more moderate separation between Colombians and Maya (0.52) (see Figure S8 for a more detailed plot).

**Figure 7.**
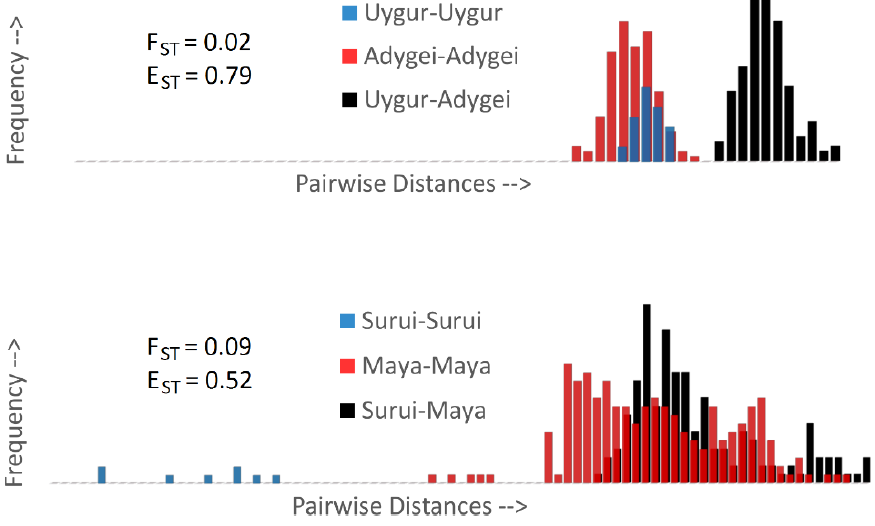
F_ST_ vs. genetic similarity in various population pairs. Pairwise distances are colored red or blue within populations and black between populations. (A) Even at a relatively low F_ST_ of 0.02 all within-population pairs among the Uygur and Adygei samples are genetically more similar than all the between-population pairs. (B) Separation is more ambiguous among Native Americans. Despite a relatively high F_ST_ of 0.09, there is substantial overlap between Maya-Maya (red) and Maya-Surui (black) samples. E_ST_ values are more consistent with the within-vs.-between population overlap and the dissimilarity fraction (ω).

## Summary and Conclusions

The core distinction between F_ST_ and E_ST_ is that F_ST_ partitions genetic diversity, whereas E_ST_ partitions genetic structure within and between populations. F_ST_ is more sensitive to differences in diversity within populations, while E_ST_ is more sensitive to outliers; though this is largely mitigated by using E_ST_median rather than E_ST_mean (see Materials and Methods). F_ST_ is weighed down by high levels of intrapopulation diversity and can be close to zero even when population clusters are completely separated. This is not necessarily a flaw in F_ST_, but it does demonstrate a conceptual discrepancy between F_ST_ and strength of clustering.

Sewall Wright proposed a series of arbitrary F_ST_ thresholds ranging from 0.05 to 0.25, denoting little to very great differentiation [1]. Notably, the highest ranking of “very great differentiation” leaves most of the range (0.25-1) undefined. Given the wider empirical range of E_ST_ and its correspondence with results from clustering schemes (Figure 2), phylogeography (Figure 5), and the dissimilarity fraction (Figure 7), such arbitrary thresholds may not be necessary for E_ST_. A value of E_ST_ larger than 0.5 simply indicates that most of the structure is between populations rather than within, corresponding to moderately separated populations such as Russians and Adygei (E_ST_=0.5), Bantu from South Africa and Kenya (E_ST_=0.48), or French and Sardinians (E_ST_=0.48) (Table S1). E_BT_, as delineated in the Materials and Methods section, performs in many ways similarly to E_ST_, though its HGDP ranking order may be intermediate between F_ST_ and E_ST_ (Table S1).

Differentiation metrics are judged by their ability to quantify meaningful evolutionary divergence, and can be indispensable in identifying *Evolutionarily Significant Units* (ESU) and *Distinct Population Segments* (DPS) for conservation [19]. For example given several subpopulations within a species, it is reasonable to prioritize the most highly differentiated subpopulation for conservation in order to maximize biodiversity. However, higher F_ST_ does not necessarily reflect stronger separation and lower misclassification, as with the Uygur and Adygei, whose clusters are better defined than those of the Surui and Maya despite a fourfold lower F_ST_ (Figure 7). In this context humans can be a useful model species simply because we know so much about human populations due to our “long habit of observing ourselves” [20]. This allows us to make educated inferences about human populations that might otherwise be overlooked, e.g., we can be skeptical of the high Surui-Karitiana F_ST_, and realize that this is most likely due to the relatively recent isolation of two small tribes. This is a luxury that we do not usually have with other species, in which case high F_ST_ can be misinterpreted as a deep phylogenetic divide, potentially leading to misguided conservation strategies. Our hope is that by combining information from both *fixation* (F_ST_) and *equidistance* (E_ST_) indices, researchers could make more informed decisions.

Unlike F_ST_, which is typically averaged across any number of markers, E_ST_ is an asymptotic measure, in the sense that it requires large datasets with many thousands of markers, which have only recently become widely accessible. With the latest SNP chips containing well over 100,000 markers, accurate estimates of departures from panmixia are finally within reach, and there is no longer a need for the simplifying assumption that subpopulations are effectively panmictic. By deriving an F_ST_-type statistic for apportioning structure within and between populations, namely E_ST_, we hope to add a new useful metric to the 21^st^ century population genetics toolkit.

## Materials and Methods

The HGDP data used in our analysis were accessed at: http://www.hagsc.org/hgdp/files.html. After removing the 163 mitochondrial SNPs and 105 samples previously inferred to be close relatives [18], the final file included 660,755 SNPs from 938 samples in 53 populations. Strings of SNPs were treated as sequences, with mismatches summed and divided by the sequence length. Pairwise distances, based on Allele Sharing Distance (ASD) [21], were calculated as one minus half the average number of shared alleles per locus. The theoretical model, mathematical proofs and numerical simulations (using Mathematica v.8.0) of SD_T_ and SD_S_ appear in Appendix A.

In the empirical analysis we used Hudson’s pairwise-distance based F_ST_ estimator [4] adapted to diploid genotypes:

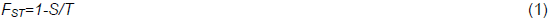

where S and T are mean pairwise distances within subpopulations and in the total pooled population.

E_ST_ was formulated in terms of standard deviations as:

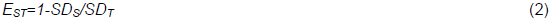

where SD_S_ and SD_T_ are the standard deviations (SD) of pairwise distances within subpopulations and in the total population. This E_ST_ estimator is referred to as E_ST_mean. We used three additional E_ST_ estimators: E_ST_min, E_ST_median, and E_ST_max (Figure S9). All four estimators use the same basic formula in (Eq. 2), with only the type of SD_S_ differing among estimators. In E_ST_min, E_ST_median, and E_ST_max, SD_S_ is respectively replaced with the smallest, median, and largest *individual SD*, where the individual SD is the standard deviation of pairwise distances between a single sample and all other samples in the population. E_ST_min uses the smallest individual SD_S_ from each population, i.e., the SD of the most panmictic sample, E_ST_median uses the median individual SD_S_, and E_ST_max uses the highest individual SD_S_. Each of these metrics has different sensitivities to various sampling biases. Due to E_ST_mean’s sensitivity to the sampling of close relatives, we used E_ST_median (which is unaffected by the inclusion of relatives as long as at least 50% of the samples are unrelated) as the primary measure of E_ST_ in this study. In the rare event that >50% of the samples are closely related, E_ST_max may be preferable, as long as at last one individual has no close relatives among the samples. E_ST_ values, especially E_ST_min and E_ST_mean, can be negative if structure is high and differentiation is low (Figure S9). Small sample sizes were often sufficient for estimating heterozygosity (Figure S10) and F_ST_ and E_ST_ (Figure S11) using all the SNPs in the HGDP dataset. Nevertheless, systematically developing estimators for E_ST_ is beyond the scope of the current treatment.

We derived an additional equidistance index, denoted E_BT_, which is less sensitive to intra-population structure and the inclusion of relatives. Recall that E_ST_ reflects equidistance (E) within subpopulations (S) compared to the total (T) population. Similarly, E_BT_ reflects equidistance (E) between subpopulations (B) compared to the total (T) population:

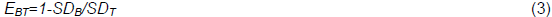

Where SD_B_ and SD_T_ are the standard deviations of pairwise distances between individuals from different subpopulations, and in the total pooled population respectively. In most cases SD_T_≥SD_B_, because SD_T_ includes pairs of individuals from the same population as well as pairs from different populations, whereas SD_B_ only includes pairs of individuals from different populations. Pairs of individuals from the same population are likely to have a higher SD due to relatives in the samples, which disrupt the panmixia (e.g., in the Naxi population, see Figures S2-S4). Panmictic populations are not just equidistant among themselves; they are also equidistant towards each other. Such populations should have similar SD_S_ and SD_B_, and thus similar E_ST_ and E_BT_. Interestingly, some East Asians populations have relatively low E_BT_, such as Cambodians vs. Mongola (E_BT_=0.13) and Japanese vs. Chinese (E_BT_=0.16). All F_ST_, E_ST_ and E_BT_ estimates in this study are based on pairwise comparisons between two populations or population groups. Each of the two paired populations was given equal weight, as were the within-and between-population pairs. Thus, 25% of the total weight was given to each population, and 50% to between-population pairs.

We developed a custom MATLAB code for extracting genetic distances from SNP data and estimating heterozygosity, pairwise distances, F_ST_, E_ST_, and E_BT_. The code corrects for missing data and small sample sizes, and identifies outliers, but includes no further assumptions or corrections. Phylogenetic trees and MDS plots were also generated with MATLAB. Equal angle and square neighbor-joining trees of individual similarities were generated from matrices of pairwise distances with the *seqneighjoin* command. An alternative script, based on the internal MATLAB *seqpdist* command for sequence distance, yielded similar results.

## Acknowledgments

We thank Alan Templeton for helpful advice, Tat Dat Tran for assisting with one of the proofs in the appendix, Lior Lesch for software support and Sagi Abelson for help with the MATLAB script.

## Appendix A *The Standard Deviation of Pairwise Distances as a Measure of Population Structure*

Our goal in this appendix is to substantiate the *asymptotic* (in terms of number of SNP loci) *standard deviation* of pairwise genetic distances as a good unsupervised measure of internal structure, thus justifiable as a basis for the definition of *E*_ST_. In particular, we prove that this asymptotic standard deviation is zero if and only if there is no internal structure (i.e., the population is panmictic).

## A model of pairwise genetic distances for genotypes from two diploid populations

Let *p_i_* denote the frequency at locus *i* of allele *‘A’* in population 1, and let and *q_i_* denote the frequency of the same allele in population 2 and assume that both populations are effectively very large and have the same contribution to the total population. The commonly-used *allele sharing distance* (ASD) measures the dissimilarity of two individual genotypes. For *diploid* genotypes, it is commonly defined as 2 minus the number of shared alleles at each locus, averaged across loci [21–22]. For multiple loci genotypes we use a normalized (by the number of considered loci) version of ASD to simplify the analysis of means and variances of the ASD distribution, as in Tal (2013). Under the assumption of Hardy-Weinberg Equilibrium, allele frequencies fully determine per-locus genotype frequencies.

Let a categorical random variable *X_i_* represent the ASD at diploid locus *i*, and let *D_n_* represent the normalized ASD across *n* loci for pairs of genotypes sampled from the *total* population,

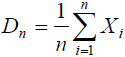

We are interested in arriving at an expression for the variance (and ultimately the asymptotic standard deviation) of *D_n_*. Under the standard assumption of *linkage equilibrium* (LE) within each of our two populations, the *X_i_* for the *total-population* pairs are *not* statistically independent, and therefore the formulation for the variance of *D_n_* requires a partition into conditional expectations. From basic principles,

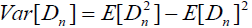

Now, to evaluate 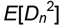 we need to condition it upon classification of pairs of genotypes as within- or between-population since the *X_i_* from the total population are not statistically independent,

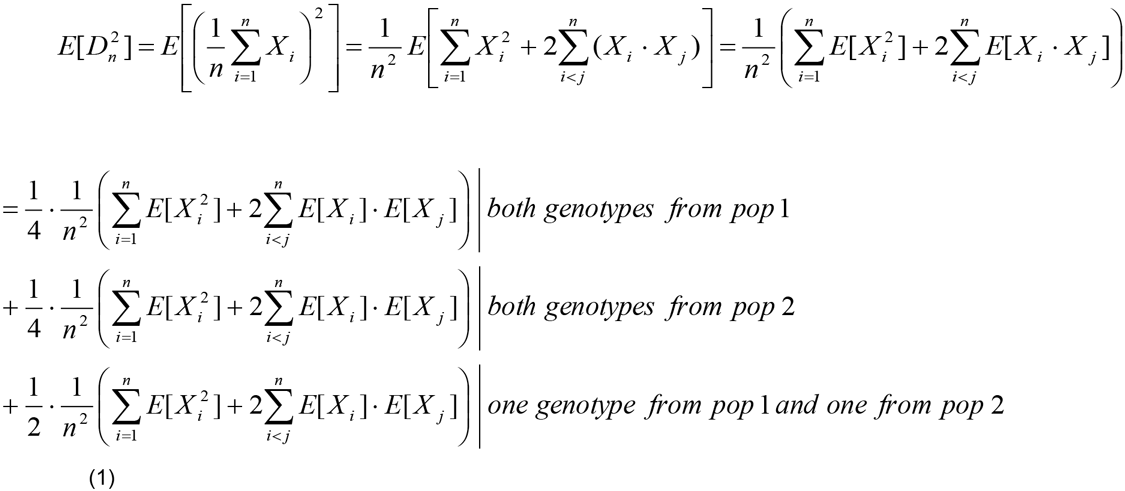

where *E*[*X_i_X_j_*]=*E*[*X_i_*]·[*X_j_*] since there is independence across any two loci for the within pairs and between pairs, and where the probabilities (assuming equal population sizes) for within-population 1 pairs, within-population 2 pairs, and between-population pairs are *at infinite population size* ¼ ¼ ½ respectively (otherwise, for finite population sizes *m* we have probabilities 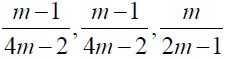 respectively).

Now, from per-locus probabilities in Tal (2013, eq. 3 and (Table 1) we derive the expected values,

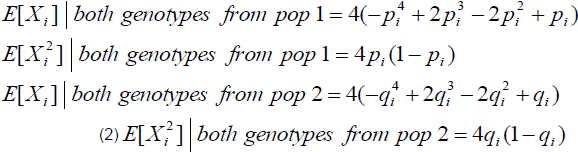

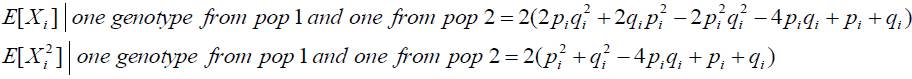

So that,

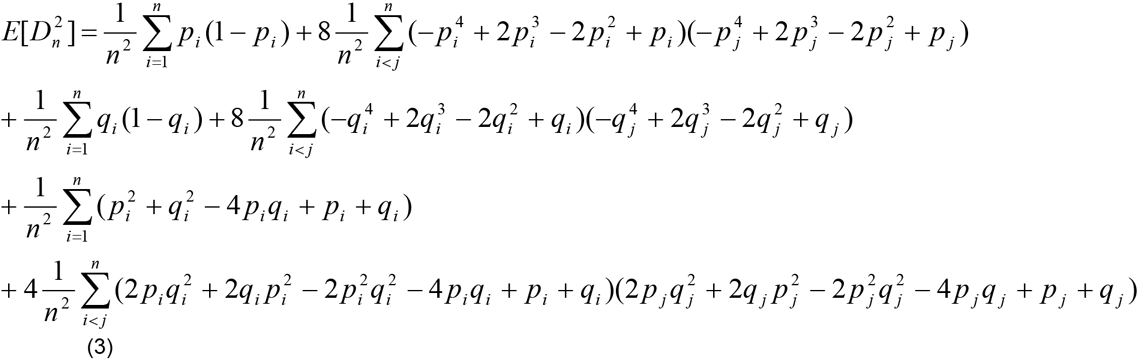

Also, since the expectation of a sum of dependent random variables is the sum of their expectations we have for the ‘total population’ *X_i_*,

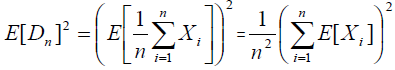

From Tal (2013, section 3.2) we have the expression for the *pmf* of *X_i_* and thus can derive E[*X_i_*],

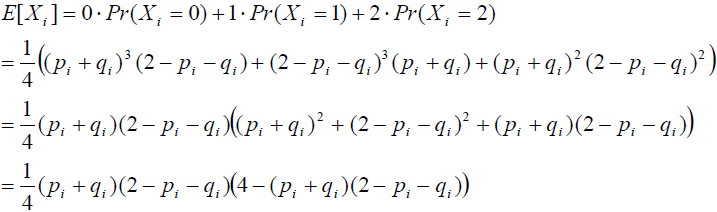

Such that,

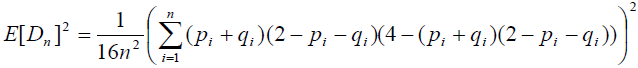

So that finally,

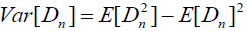

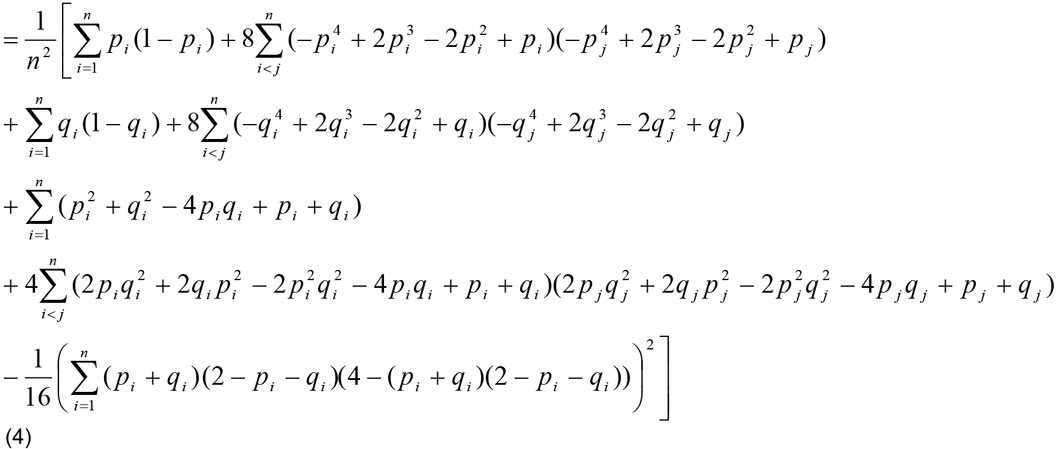

Thus we have an explicit formulation for the variance of the pairwise distance distribution of genotypes from two panmictic populations in terms of the allele frequencies across a given number of loci, *n*.

Crucially, we would like to prove that at the limit, the pairwise distance variance is asymptotically above *zero if and only if* the population has internal structure; i.e., if for any *F_ST_*>0,

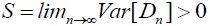

We will proceed by deriving an explicit expression for *S*. Consider an *equivalent setting* comprised of three random variables *W,Y* and *Z*, which represent the pairwise distances of genotypes within population 1, within population 2 and between populations 1 and 2, respectively. We sample *n* values *X_i_* from just *one* of these distributions, by first flipping a 3-sided coin to decide from which: with a probability α for *W*, a probability *β* for *Y* and a probability γ for Z. Once the distribution has been selected, the sampling of *X_i_* is done *i.i.d*. Note that due to the randomized choice of the distribution from which to sample *all* the *X_i_*, they are identically distributed but *not* independent. Now we set,

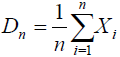

We would like to get an expression for *S* in terms of the expectations of *W, Y, Z* and α, β, γ, where,

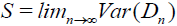

From the law of total variance,

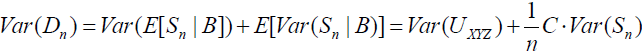

where *B* is a categorical random variable that describes which of the distributions *W*, *Y*, *Z* we are sampling from, with probabilities α, β, γ respectively, and where *U_XYZ_* is a discrete random variable taking the values of *μ_W_*=*E*[*W*], *μ_Y_*=*E*[*Y*], *μ_z_*=*E*[*Z*] with corresponding probabilities α, β, γ respectively. Hence at the limit n→∞ we have,

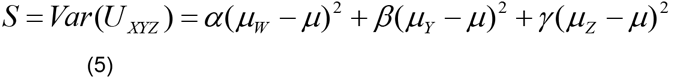

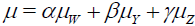

and *S*=0 *if and only if* the three means are equal, i.e., *μ_W_*=*μ_y_*=*μ_z._*

Now consider three *sequences* of random variables *W_i_*, *Y_i_*, *Z_i_, i:1,…,n*, instead of the three single random variables, and sample *n* values from one of these sequences (again according to the prior probabilities α, β, γ). Once the sequence is selected, these samples are independent but now *not identically distributed*. We would again like to find *S*, and more importantly, the condition for which it is zero, this time in terms of *E*[*W_i_*], *E*[*Y_i_*], *E*[*Z_i_*](and the prior probabilities). Sampling from a sequence with fixed probabilities just defines a new mixture distribution −s o the problem gets reduced to the one already solved. Therefore *U_XYZ_* is now defined by the three limits (since we have derived *S* in Eq. (5) at the limit n→∞),

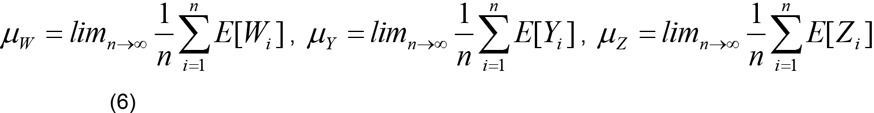

with probabilities α, *β*, γ.

Crucially, this sampling scenario corresponds to our original setting of formulating the variance of the genetic distance of genotypes sampled from the total population, given the sequencing of an infinite number of loci, where *μ_W_* and *μ_Y_* in Eq. (6) represent the two within-population pairwise distance means and *μ_Z_* the total-population mean (derived below), and where the respective probabilities are as in Eq. (1), α=¼, *β*=¼, γ=½assuming *infinite population size*. Again, S=0 *if and only if* these means are equal, i.e., *μ_W_*=*μ_y_*=*μ_y_*. Let us analyze the conditions for these equalities, given the corresponding formulations of the pairwise distance means. First, using the additivity of expectations,

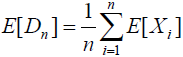

we get from Eq. (2) the expressions for any *finite n*,

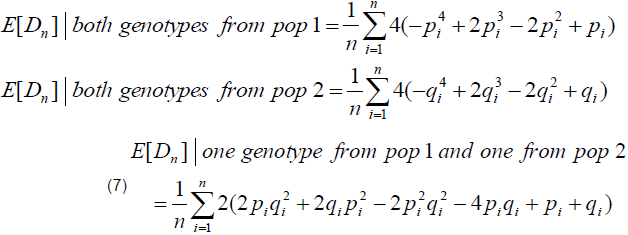

bearing in mind the analysis pertains to *E*[*D_n_*] as *n*→∞. We proceed to examine what can be concluded from the equalities *μ_w_=μ_Y_=μ_z_* (the only case where *S*=0), given the means in Eq. (7), about the allele frequencies *p_i_* and *q_i_* for any finite *n* (and this also holds at *n*→∞). Thus we start by explicitly writing the equalities (where the 1/*n* cancels out),

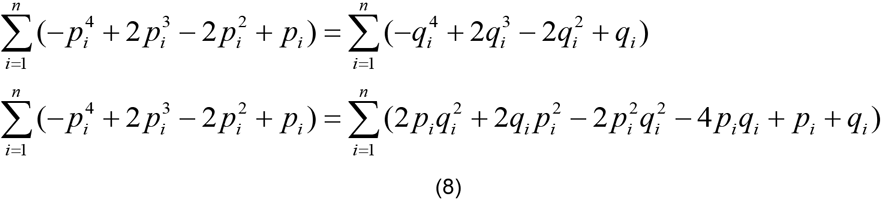

To proceed we substitute new variables,

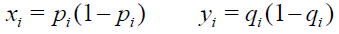

Then, the 1st equation in (8) becomes,

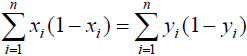

such that,

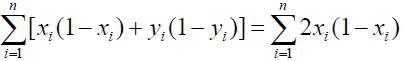

and using the 2nd equation in (8),

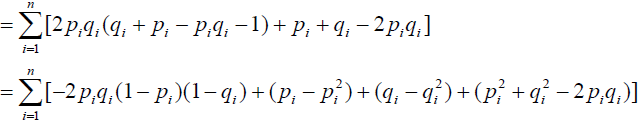

And again in terms of the new variables,

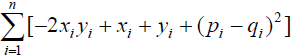

This implies that,

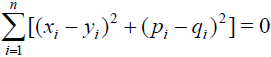

which occurs only if *p_i_* = *q_i_* for all *i* = 1,…,*n*.

Therefore the *asymptotic* variance of the pairwise genetic distances (normalized by number of loci) of genotypes sampled from the combined population, comprising two subpopulations, is zero *if* this combined population is essentially a single panmictic population (i.e., *p_i_*=*q_i_* for all *i*, or *F*_ST_=0). Since we have defined *E_ST_* in terms of standard deviations rather than variances, we will subsequently consider the *asymptotic standard deviation *SD*_T_*, which is simply defined as the square root of the *asymptotic variance, S* for the ‘total’ population. Figure A1 depicts numerical simulations of both *SD_T_* and the average within-population SD (*SD*_S_) for our two population model, as a function of the number of SNPs considered. While *SD_S_* converges to zero, *S_DT_* asymptotes to a value greater than zero, revealing the underlying structure. We note here that the rate of convergence to zero for *SD_S_* is highly dependent on the diversity of each population – for lower diversity the within-population SD converges faster (and thus tends to be lower for any finite number of loci). This factor also influences the rate of convergence of *E*_ST_ to its asymptotic value (see main text, Figure 3).

**Figure A1.**
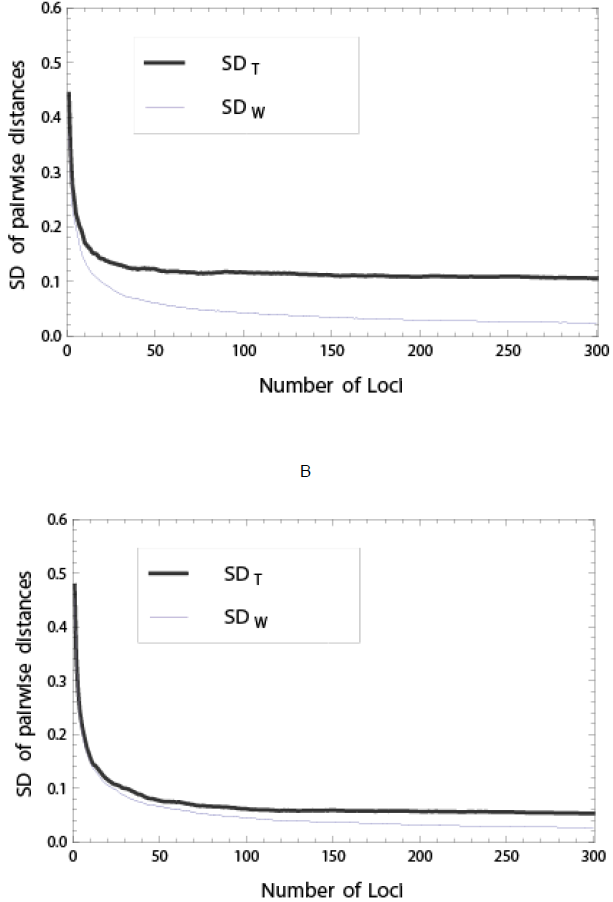
A simulation of *SD*_T_ and *SD*_S_ under a two panmictic population model demonstrating the divergent behavior of these two statistics with an increasing number of SNP loci. SNP frequencies are modeled on Beta distributions (as in [5]). **A**: with *F*_ST_=0.10. **B**: with *F*_ST_=0.03.

To further substantiate *SD*_T_ as a measure of structure, we would like to characterize the relation of *SD*_T_ to *F*_ST_, both formulated as expressions of allele frequencies from two populations. We will proceed numerically, as our goal here is merely to get a qualitative intuition into the association of the two statistics.

We have from Eqs. (5), Eq. (6), Eq. (7), that asymptotically as *n*→∞, or practically under a high number of SNP loci,

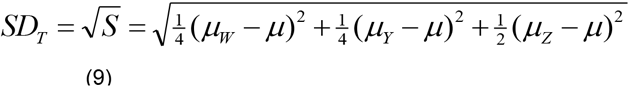

where,

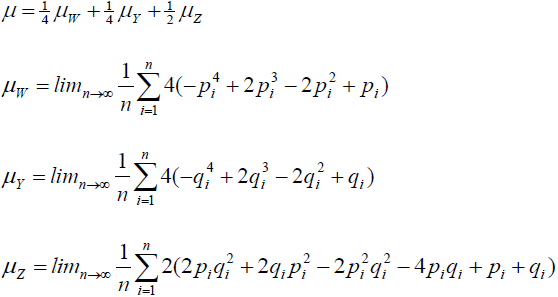

And from Tal (2013, Eq. 10) we use the most common expression for *F*_ST_ across any number of *n* SNPs,

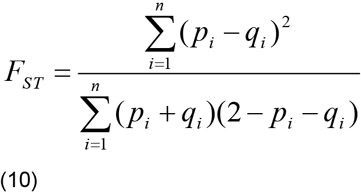

Under the standard assumption that SNP frequencies are modeled on a *Beta* distribution with parameters deriving from some historical process (see [5, 22]) we sample a large number of sets of SNP frequencies for two populations, each set generated from two *Beta* distributions with some randomized parameters. For each set we compute the pair *SD*_T_ (Eq. 9) and *F*_ST_ (Eq. 10) to generate a scatter plot of their association. Figure A2A-D depicts several typical instances of such a simulation, demonstrating that the correlation of the two statistics is substantial in the case of two or more panmictic populations,

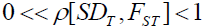

and is lower if the subpopulations are structured. Extending the numerical analysis for any number of *panmictic* multiple populations indicates that *SD*_T_ closely follows *F*_ST_ as a measure of structure.

**Figure A2.**
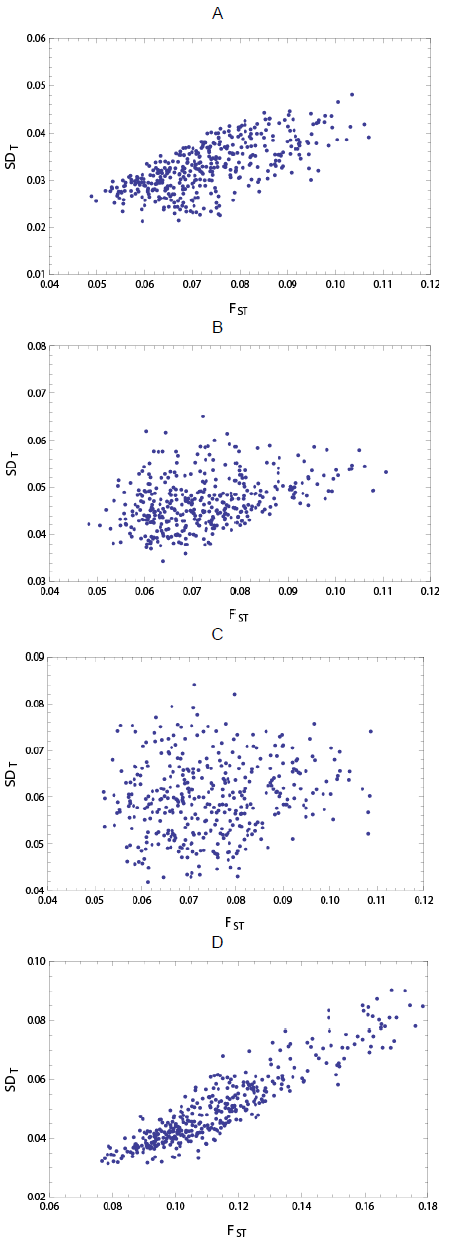
Scatter plots indicating a positive correlation for *SD*_T_ and *F*_ST_. Each dot represents the two statistics computed for data sampled from our population model with 1000 SNPs and allele frequencies from Beta distributions. The Pearson product-moment correlation coefficient of *F*_ST_ and *SD*_T_ is 0.67 for plot **A** with panmictic populations, 0.38 for plot **B** with low structure populations, 0.14 for plot **C** with high structure populations, and for 0.94 for plot **D** with three panmictic populations.

We may generalize this result to more than two populations in a straightforward manner using induction. For instance, for three populations Eq. (9) becomes,

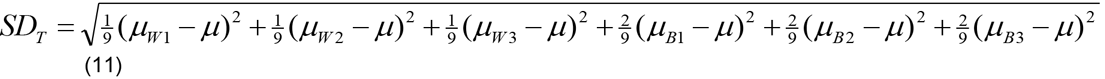

where,

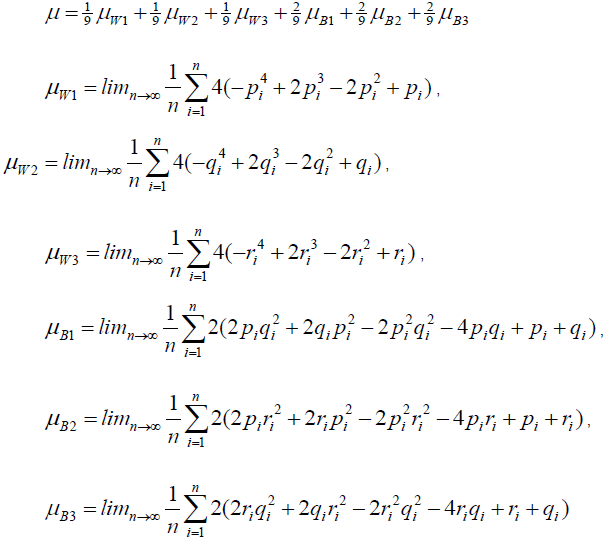

and where *F*_ST_ for 3 panmictic populations with SNP loci is derived in the same manner as Eq. (10),

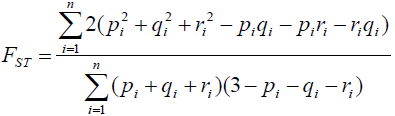

A further perspective into *SD*_T_ as an unsupervised measure of internal structure is afforded by a qualitative comparison with principal component analysis (PCA) plots on data generated by the model. PCA is an *unsupervised* technique, essentially a dimensionality reduction procedure, used to emphasize the directions of greatest variation and bring out any strong patterns in a dataset. It can be used as a ‘preprocessing’ stage for clustering high-dimensional data, such as characteristic of population genetic samples. In such a setting, the first principal components tend to also extract existing substructure within the data in the form of clusters [14]. But more crucially to our goals, the relative dispersion of clusters on a PCA plot is highly associated with their internal structure, i.e., departures from panmixia, with increasing number of loci (and asymptotically, panmictic clusters would diminish to a single dot). This property is congruent with the convergence of *SD*_T_ to some value strictly greater than zero for non-panmictic populations. This is depicted in the four PCA plots of the same populations under increasing SNP count in Figure A3A-D.

**Figure A3.**
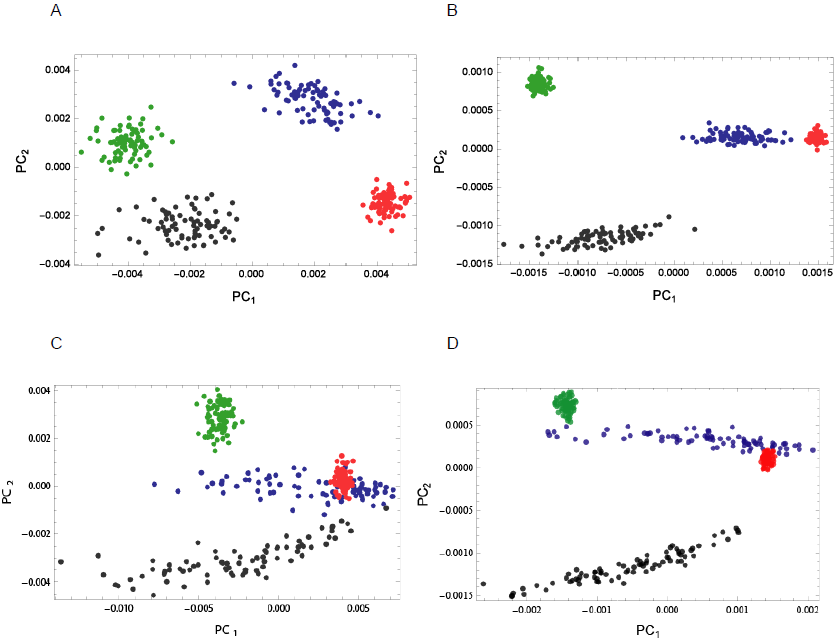
PCA plots from simulated SNP data of four populations (80 samples each) demonstrating the much pronounced decrease in *SD*_S_ for panmictic populations (red and green) relative to a structured ones (black and blue), for two different patterns of internal structure, as the number of SNP loci processed by the PCA scheme is increased. **A-B**: from 1K SNPs to 8K SNPs, where structure results from some random LD pattern. **C-D**: from 1K SNPs to 8K SNPs, where structure results from an LD pattern resembling admixture.

Finally, through numerical simulation of our model, we can see how varying degrees of internal structure (simulated by controlling the LD patterns) result in different asymptotic levels of *E*_ST_ (Figure A4). This serves to substantiate the empirical analysis depicted in **Figure 3** of the main text.

**Figure A4.**
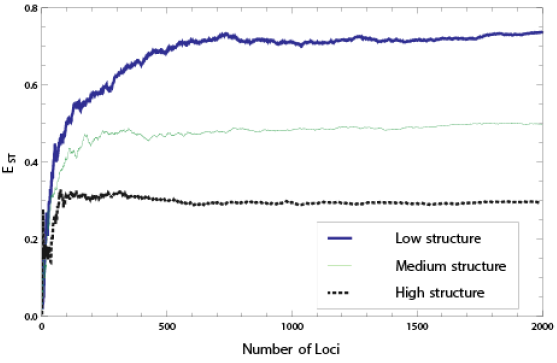
A numerical simulation of a model for *E*_ST_ for two structured populations (with *F*_ST_=0.05). *E*_ST_ was computed using the formulation in Eq. (2) of *Materials and Methods*.

